# Proximity to Visual Stimuli Reduces Post-Saccadic Alpha Lateralisation

**DOI:** 10.1101/2024.10.09.617405

**Authors:** Christopher Turner, Aleksandra Vuckovic, Gemma Learmonth, Alessio Fracasso

## Abstract

Posterior alpha lateralisation is a well-established marker of visual spatial attention, with growing evidence suggesting that it may also be linked to oculomotor planning and execution. However, saccades also result in a change in visual input to the fovea, which could be linked to lateralised posterior alpha. In this study, we investigate the interplay between saccade-induced foveal input changes and lateralised posterior alpha by analysing saccades from the cue period of a Posner task and the active feedback period of a neurofeedback task, where visual stimuli were present on screen. Specifically, we examine the impact of saccade landing proximity to visual stimuli and saccade amplitude on post-saccade alpha lateralisation. Consistent with previous research, we observe a general post-saccade alpha lateralisation across both tasks. Importantly, we find that alpha lateralisation is reduced in the neurofeedback task when saccades land close to visual stimuli compared to those landing further away, while saccade amplitude has no significant effect. This investigation highlights the importance of controlling for both eye movements and visual stimuli when investigating alpha lateralisation in fixational tasks. Moreover, it shows how both oculomotor and afferent processing mechanisms affect posterior alpha lateralisation.

## 1 Introduction

Numerous studies have demonstrated that posterior EEG power within the typical 8-12Hz alpha band is lateralised during spatial attention tasks. This lateralisation is characterised by increased alpha power in the posterior cortex ipsilateral to the attended direction, and decreased alpha power in the contralateral hemisphere (Kornrumpf et al., 2017; Peylo et al., 2021; Rihs et al., 2007; Sauseng et al., 2005; Thut et al., 2006; Worden et al., 2000). Lab-based studies often require participants to maintain gaze fixation towards a central fixation point to control for overt eye movements which can introduce EEG artefacts (e.g. Coldea et al., 2021; Croft & Barry, 2000; Desantis et al., 2020; Rihs et al., 2007; Sauseng et al., 2005; Thut et al., 2006). However, this overlooks the fact that small, fixational eye movements such as microsaccades are always present during fixation (Engbert & Kliegl, 2004; Martinez-Conde et al., 2004).

Recently, interest has grown in how saccades contribute to lateralised alpha power, linking eye movement planning and execution to EEG alpha band modulations (Balestrieri et al., 2024; Liu et al., 2022, 2023; Popov et al., 2021). For example, Popov et al., 2021 report alpha modulations in the presence of saccades, even when visual input is absent. Further, exogenous attention experiments report modulation of alpha power after displaying unpredictable cues which cannot be anticipated by saccades (Balestrieri et al., 2024; Keefe & Störmer, 2021; Landry et al., 2024). Balestrieri et al., 2024 argue that these alpha modulations are a product of the oculomotor system through a suppression of saccades towards the exogenous cue.

In further support of the oculomotor influence on alpha lateralisation, Liu et al., 2023 demonstrated that alpha-band EEG signals can be modulated by saccades during a delay period in a working-memory task when only a fixation cross is present on the screen, reinforcing the link between alpha power and eye movement planning and execution. However, the interaction between visual input (visual afferent) and alpha modulations *following* an eye movement has not been investigated so far.

Visual stimuli have been shown to directly modulate the EEG signal. For example, changing visual stimuli on the fovea can modulate the EEG activity broadly (Nathan & Hanley, 1975); alpha oscillations and lateralisation can be induced by visual stimulation (Schürmann & Başar, 2001); and alpha oscillations can be induced by new visual stimuli (Balestrieri et al., 2024; Landry et al., 2024; Vanni et al., 1997). Since eye movements alter visual input on the fovea, it remains unclear the extent to which post-saccade alpha modulations are influenced by oculomotor planning/execution and visual changes due to the eye movement itself (i.e. changes in visual afference). Understanding this is crucial for designing and interpreting EEG experiments requiring stable fixation because EEG modulations may result from both oculomotor planning/execution and visual afference. Furthermore, in this investigation we can uncover whether the saccade-locked alpha lateralisation signal is predominantly linked to mechanisms of oculomotor planning/execution or visual afference. In the former case, the saccade-locked alpha lateralisation signal could be interpreted as a correlate of covert spatial attention (Kornrumpf et al., 2017; Peylo et al., 2021; Rihs et al., 2007; Sauseng et al., 2005; Thut et al., 2006; Worden et al., 2000), or as a corollary to the saccade motor signal (Crapse & Sommer, 2008). In the latter case, the signal could be associated to the retinal displacement of the visual stimulus while the eyes are moving, or to the influence of immediate post-saccadic visual afference (Kagan et al., 2008; Snodderly, 2016).

Here we present an analysis of simultaneous eye-tracking and EEG data that were recorded during a covert visual-spatial attention-based EEG neurofeedback experiment. Participants completed three sessions of combined transcranial alternating current (tACS) and EEG neurofeedback in a within-subjects design. The presence of lateralised visual stimuli during the tasks allowed us to investigate the interaction between oculomotor planning/execution and visual afference. In line with Liu et al., 2023, we expected to observe a transient alpha band lateralisation after saccade onset. We extend this by investigating whether and how the EEG signal is modulated by the post-saccadic visual afference; testing whether alpha lateralisation is modulated by the proximity of the saccade end point relative to the on-screen visual stimuli.

## 2 Methods

### 2.1 Participants

Thirty healthy participants aged 16 and over with normal, or corrected to normal, vision participated in this study (mean age = 27.26 ± 8.98, 24 female, 29 right-handed). Data from four participants were not included in the analysis due to three withdrawals after the first session because of scheduling conflicts, and one from difficulties recording eye tracking data through the participant’s glasses. Further, one session from two participants were lost due to an issue saving the data. In total, 24 full datasets and two partial datasets were used in the final analysis. All participants provided written, informed consent prior to participation, and were offered £40GBP after completion of all three sessions. This study was approved by the University of Glasgow Medical, Veterinary & Life Sciences ethics committee and was performed in accordance with the Declaration of Helsinki.

### 2.2 Task and procedure

Each participant attended three sessions on different days, where they completed the following three conditions in a counterbalanced order: 1) sham neurofeedback and alpha tACS, 2) active neurofeedback and alpha tACS, and 3) active neurofeedback and gamma (40Hz) tACS.

#### 2.2.1 Posner task

Manual reaction times were measured with two Posner tasks within each session; one at the beginning and one at the end. Each Posner task had four equal blocks of 25 trials totalling 200 trials per session. **Figure 1a** shows the individual trial structure which consisted of a fixation period of 1.5s followed by a cue period where a target appeared after a pseudo-random delay between 2.5s and 4s after cue onset, and finally a response period of 1s. During the fixation period participants were instructed to maintain fixation on a central point. Two lateralised blue circles (3.5° diameter, vertically offset by −1° and horizontally offset by +/− 5° visual angle), one to the left and one to the right of the central fixation point, were also present on the screen. During the cue period, a directional arrow appeared behind the fixation dot indicating the direction that the target was likely to appear. This cue had three possible orientations with equal probability of occurring: left, right, or both directions. After the pseudo-random delay, a target would appear either on the same side as the cued direction 70% of the time (valid trials) and 30% on the opposite side (invalid trials). Once the target appeared, the participant was instructed to press the arrow key on a keyboard placed in front of them corresponding to the side on which they perceived the target. Participants used the index and middle fingers of their dominant hand to respond to the left and right targets. For the analysis presented here, we focus on the cue period of the Posner task. Saccades were taken from 250ms after the cue onset until 2100ms after cue onset to avoid cue-related ERPs in the signal and to include the maximum length of the cue period before a possible target appeared. This enabled us to investigate the effects of visual stimuli on any saccade-induced EEG signal while participants were fixating.

**Figure 1:**
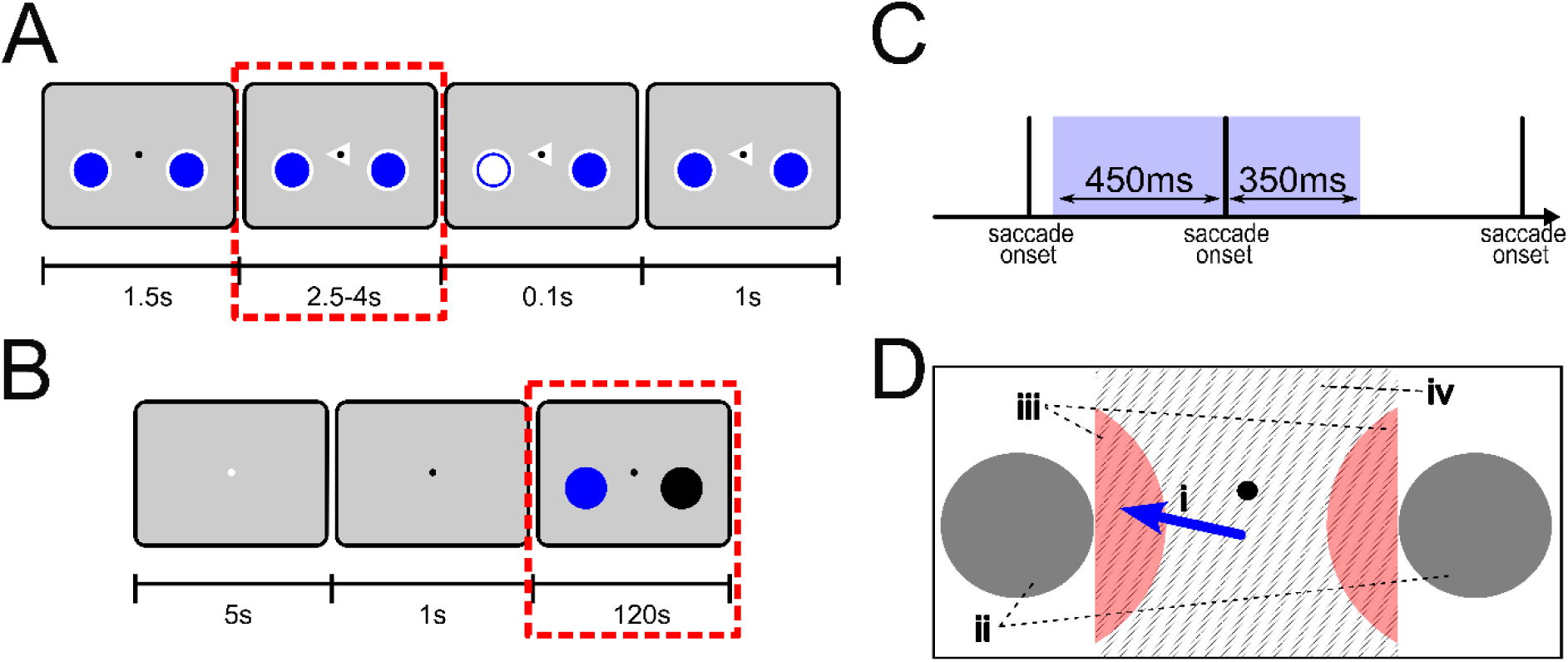
Task structure, saccade detection and classification. **A**) Posner task trial structure with saccades taken from the cue period outlined by the dashed red line. **B)** Neurofeedback task structure with saccades taken from the active neurofeedback period outlined by dashed red line. **C)** Temporal threshold surrounding saccades. Only saccades that occurred in isolation (i.e. with no previous or subsequent saccades occurring within the blue shaded area) were included in the analysis. **D)** Saccade proximity classification. Saccade proximity was calculated from the landing point of each saccade indicated by the end of the blue arrow (i) to the centre of the lateralised visual stimuli (ii). The median saccade proximity for each participant was calculated over all sessions. This was used to determine the proximity threshold indicated by the red shaded region (iii). Saccades ending within this threshold were classified ‘close’ and those ending outside this threshold were classified as ‘far’. In the example, the saccade would be a ‘close’ saccade as it ends within this threshold. Only saccades that began and ended between the inner boundaries of the lateralised visual stimuli were included. Boundaries indicated by the grey shaded area (iv).

#### 2.2.2 Neurofeedback task

During each session participants completed four blocks of three neurofeedback trials totalling 12 trials per session. **Figure 1b** outlines the trial structure. Each trial began with a five second rest period followed by a one second fixation period indicating that the participant should prepare for the neurofeedback section to start. Two minutes of active neurofeedback followed, where the participants attempted to modulate the left of two horizontally lateralised circles. These lateralised targets were in the same position on the screen as the lateralised targets in the Posner task (3.5° diameter, vertically offset by −1° and horizontally offset by +/− 5° visual angle). During the neurofeedback training period, participants were instructed to maintain their fixation on the central fixation point and modulate their covert visuo-spatial attention to focus on the left target. They were told that this would allow them to change the target colour from red to blue and that they should attempt to make the target as blue as possible for as long as possible. However, participants were also advised that they could use their own strategy if they discovered one that works for them.

During the active neurofeedback sessions, alpha power at individual alpha frequencies (IAF) modulated the colour of the left visual stimulus with the right target remaining black. Specifically, the colour was modulated by the participant’s alpha asymmetry index (AAI) from the P5 and P6 electrodes. The AAI was calculated as (αP5-αP6)/(αP5+αP6) where αP5 and αP6 are the alpha power from the P5 and P6 electrodes respectively. Alpha power in the IAF range was calculated using the *FFT envelope detection* option with a 1000 sample window size in the NFBLab software (Smetanin et al., 2018) with an additional 100-sample rolling filter to smooth the feedback signal. The IAF was derived as the peak alpha frequency +/− 2Hz and was calculated from the initial eyes-closed resting state EEG recording for each session using the *savgol_iaf* function from the Philistine python package (Alday et al., 2023), which is based on Corcoran et al., 2018.

For the current analysis, saccades were extracted from a period starting five seconds after the start of neurofeedback to the end of the neurofeedback period. This was to avoid visually evoked ERPs due to the onset of the neurofeedback period which, due to software, could potentially be delayed.

### 2.3 EEG data acquisition

A BrainAmp MR amplifier (Brain Products GmbH, Germany) with a 64-channel passive electrode BrainCap were used to record EEG data at a sampling rate of 1000Hz. Electrodes were placed according to the 10-10 international system with the ground at AFz, online reference at FCz, and one additional electrode placed on the right outer canthus to record eye movements. Electrodes O1 and Cz were removed from the cap due to tACS electrodes placed at these locations. BrainVision Recorder software was used to record EEG data during the Posner task, and a modified version of the NFBLab neurofeedback software (Smetanin et al., 2018) was used to record EEG data during the neurofeedback task.

### 2.4 Eye tracking

Participants were seated with their heads resting on a chin rest 60cm in front of the display monitor. An SR Research EYElink1000 was used to record binocular eye tracking data with a sampling rate of 500Hz. At the beginning of each Posner task, and each block of neurofeedback trials, a five-point calibration routine was performed. Drift corrections were performed at the beginning of each block of the Posner task.

### 2.5 Saccade Detection and Classification

Binocular eye tracking data was interpolated to 1000Hz. An algorithm adapted from (Nyström & Holmqvist, 2010) was then used to detect saccades from the interpolated data (Niehorster et al., 2015). This algorithm uses adaptive thresholds to detect saccades based on peak velocity, and duration while also separating glissades (i.e. a small wobble of the eye after a saccade that is often not accounted for in event detection algorithms). A further conjugate eye movement check was then run to separate saccades from noise. An eye movement was determined to be conjugate if an eye movement could be detected from the left and right eye within a 10ms time window of each other.

A time threshold of 450ms before and 350ms after each saccade was used to ensure observed effects in the EEG signal was due to individual saccades (i.e. only saccades that occurred in isolation were included in the analysis, see **Figure 1c**). Leftward and rightward saccades were classified by calculating the difference between their end and start positions (end horizontal position minus start horizontal position) in the x-y screen coordinate system. In this manner, leftward saccades were characterized by a negative sign, and rightward saccades by positive difference. Leftward and rightward saccades were then classified further based on their end position proximity to the lateralised visual stimuli, and on the amplitude of the saccade. For the proximity classification, the Euclidian distance was calculated between the centre of the left and right lateralised targets, and the end x-y coordinate of each saccade for each participant. The median of these distances was then used to define a threshold region to separate ‘close’ (saccades closer to the lateralised target) and ‘far’ (saccades further from the lateralised target) per participant (**Figure 1d**). For the amplitude classification, the Euclidian distance between each saccade’s start and end x-y coordinates was calculated per participant (i.e., saccade amplitude). The median amplitude was used to separate ‘small’ and ‘large’ saccades per participant. Only saccades with end coordinates within the bounds of the inner radius of the lateralised targets were included. This was to ensure lateralised targets started and finished on the same side of each saccade (**Figure 1d**).

To remove extreme values outside the range of normal eye movements, saccades with a peak velocity greater than 650deg/sec were removed. Further, saccades with an amplitude less than 1arcmin were removed. Following these criteria, on average 3.3 (SD = 2.9) saccades were removed per participant.

### 2.6 EEG Preprocessing

The EEG data was pre-processed with the MNE Python software v1.4.2 (Larson et al., 2023). Eye blinks, large lateral eye movements, and muscle artefacts were cleaned from the raw EEG data for each session using Independent Component Analysis (ICA). This process involved first filtering the data between 1 and 40Hz and removing bad channels. The ICA algorithm was run over this filtered data using the *fastica* method in the MNE Python ICA class. Bad ICA components were identified by manual visual inspection and were then removed from the raw data.

To examine the impact of eye movements on the EEG data, the ICA-cleaned, raw EEG data were filtered between 0.1 and 40Hz and bad channels interpolated. We then extracted epochs 1000ms around each saccade onset and down sampled to the data to 250Hz. Any epoch with an instantaneous voltage greater than 150uV in any channel was removed.

### 2.7 Saccade Parameters

We visualised the main sequence of each participant in each task (saccade peak velocity as a function of saccade amplitude). Specifically, we took the quantile median of 10 quantiles for each participant for saccade amplitude and peak velocity. Further, we visualise the temporal distribution of saccades within each task by displaying box plots for 23×5000ms bins in the neurofeedback task, and 18×75ms bins in the Posner task. Finally, we visualise the spatial distribution of saccades in both the Posner and Neurofeedback tasks by plotting the average saccade direction in 90 bins on a polar plot across participants.

### 2.8 Statistical Analysis

The epoched time-series EEG data was converted to a time-frequency representation using Morlet wavelets with the *tfr_morlet* function in MNE Python v1.4.2. Spectral power was calculated between 2Hz and 40Hz in 1Hz steps with the number of cycles varying at 50% of the frequency using fft based convolution. The time-frequency decomposition was averaged over all participants for each saccade group (i.e., *small-close, small-far, large-close, large-far*) and direction (*leftward, rightward*) and cropped to −250ms and 400ms around saccade onset.

We chose to conduct a similar analysis to Liu et al., 2023. This allowed us to determine whether ***i***) we could observe the saccade-locked power modulation within the 8-12Hz alpha band, and ***ii***) we could test whether this could be modulated by the proximity to external visual stimuli or by saccade amplitude. Similar to Liu et al., 2023, a lateralisation contrast was calculated with the time-frequency data from posterior electrodes PO7 and PO8 contralateral and ipsilateral to saccade direction (i.e. ((contra-ipsi)/(contra+ipsi))x100). The grand average of this lateralisation contrast was calculated across all participants and investigated with a two-tailed, non-parametric cluster-level paired t-test. This was implemented with the *permutation_cluster_1samp_test* function in MNE Python with 10000 permutations and a cluster forming threshold corresponding to a p-value of 0.05. Clusters were determined to be significant if the cluster p-values were less than 0.05. We visualised the whole scalp topography of the alpha band by subtracting the leftwards and rightwards saccades (specifically ((leftwards-rightwards)/(leftwards+rightwards))x100). Grand average topography maps were baselined from −400ms to −200ms.

To examine the role of saccade amplitude and proximity to external visual stimuli on the saccade-induced EEG modulation, we evaluated differences across saccade groups in the time-frequency lateralisation contrasts using a cluster-based permutation approach. First, time-frequency lateralisation contrast data were baselined by subtracting the mean of a baseline window from −250 to −50ms before saccade onset. Then, the *permutation_cluster_test* function in MNE Python was employed with 10000 permutations and a cluster forming threshold corresponding to a p-value of 0.05. A two-tailed 2×2 repeated measures ANOVA was used as the statistical function. This was implemented in the *f_threshold_mway_rm* function from the stats package in the MNE python library. The ANOVA factors were saccade proximity (levels: close and far), and saccade amplitude (levels: large and small). Clusters were determined to be significant if the cluster p-values were less than 0.05.

## 3 Results

In total, 9691 (M=372.73, SD=180.45) saccades were detected for the Posner task and 29884 (M=1149.38, SD=550.88) saccades were detected for the neurofeedback task across participants. Saccades show a close relationship between peak velocity and amplitude, following the typical main sequence (**Figure 2a, b**) (Bahill et al., 1975; Zuber et al., 1965). **Figure 2c** and **Figure 2d** show the temporal distribution of saccades across trials in the Posner and neurofeedback tasks respectively. Saccades are evenly distributed over both tasks with a larger distribution of saccades from 450ms to 600ms after the onset of the cue in the Posner task. **Figure 2e** and **Figure 2f** show saccades are mostly horizontally distributed, followed by saccades distributed in the upwards (positive 90°) direction for both the Posner and neurofeedback task. Individual participant saccade distributions are similar across participants. A similar number of saccades were detected for both leftward and rightward directions within each task allowing for a comparison between groups (**Table 1**).

**Figure 2:**
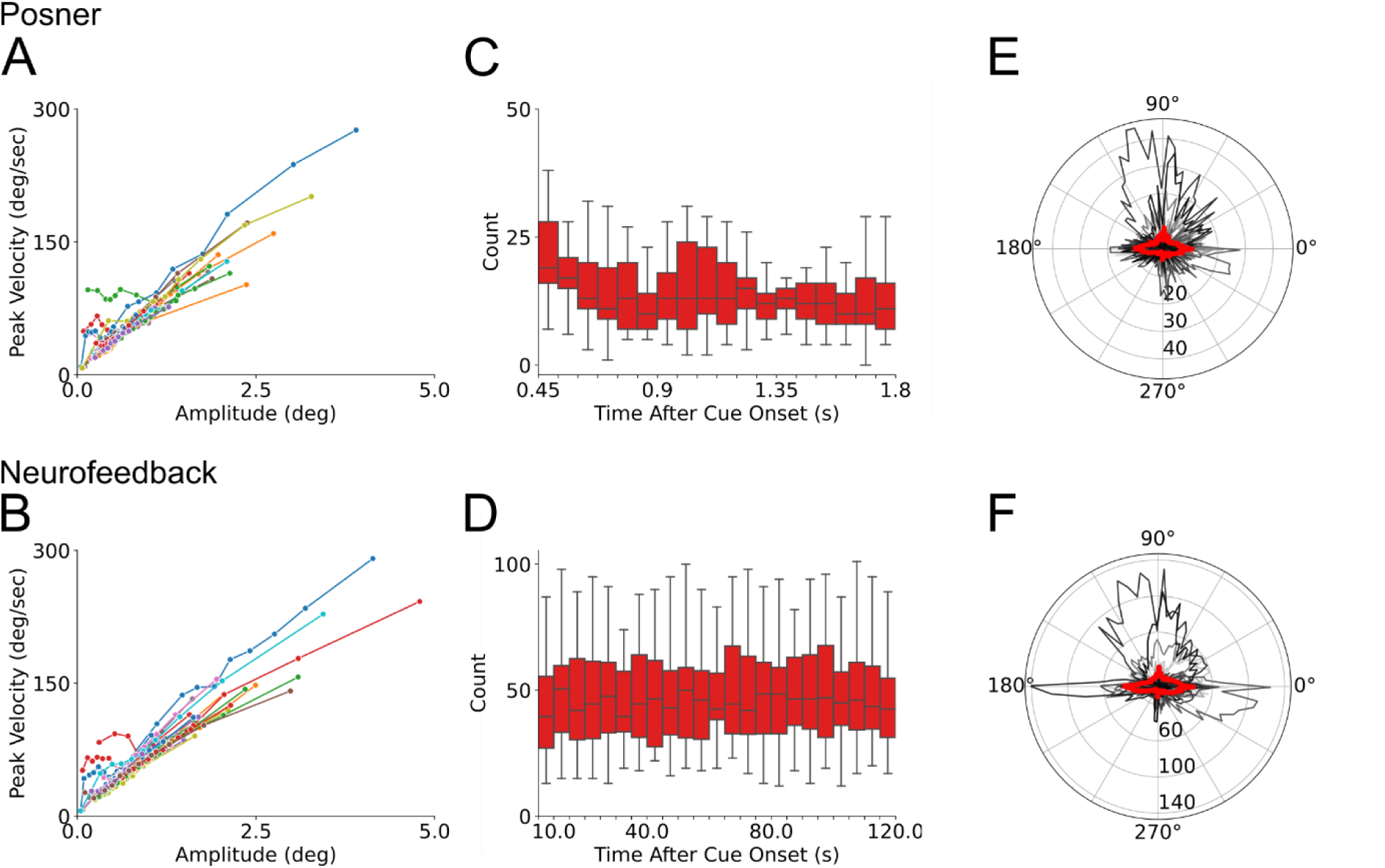
Plots illustrating different parameters of the measured saccades for the neurofeedback and Posner task. **A**) Main sequence (saccade peak velocity as a function of saccade amplitude) per participant for the Posner task. Each coloured line shows the quantile median over 10% quantiles for each participant. **B**) Main sequence per participant for the neurofeedback task **C)** Box plots showing the number of saccades across the length of a single trial for 18 bins of 75ms duration for the Posner task. **D)** Box plots showing the number of saccades across the length of a single trial for 23 bins of 5000ms duration for the Neurofeedback task **E)** Posner task polar histogram from 90 bins showing saccade distribution for all saccades. Each grey trace represents a separate participant with the red trace representing the average over all participants. **F)** Neurofeedback task polar histogram from 90 bins showing saccade distribution for all saccades

**Table 1.**
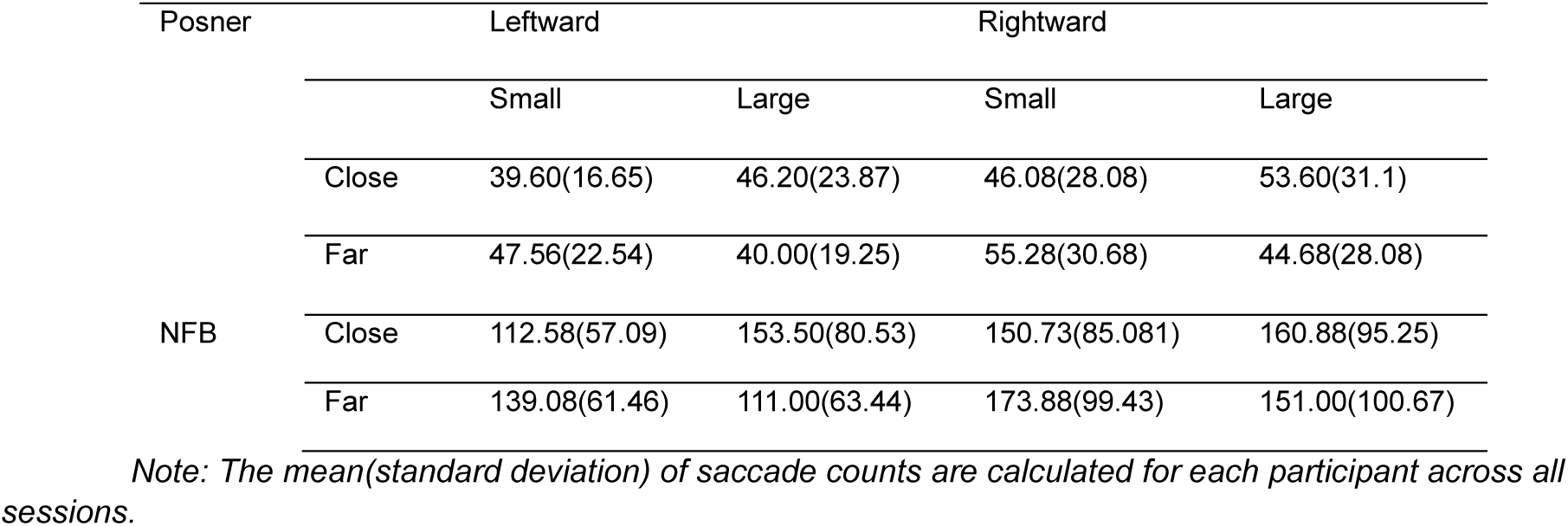
Posner and neurofeedback (NFB) task saccade averages.

### 3.1 Transient Lateralisation of EEG alpha

The difference in EEG activity from the posterior electrodes (PO7 and PO8) contralateral and ipsilateral to the saccade direction shows a clear lateralisation around the alpha band. This can be seen by the negative difference in power values after the onset of the saccade in **Figure 3c**, and **Figure 3d**. A non-parametric cluster-level paired t-test shows significant clusters for the Posner task (cluster p-value = 0.003) and the neurofeedback task (cluster p-value < 0.001). Significant clusters for both the Posner and neurofeedback tasks are at similar frequencies and similar times (lasting until approximately 250ms after saccade onset). The transient nature of the alpha band lateralisation can be further seen in **Figure 3e** and **Figure 3f**, beginning around the saccade onset and extending until approximately 250ms after saccade onset for the Posner task, and 300ms after saccade onset for the neurofeedback task. As the above-mentioned results are a relative difference in alpha power between contralateral and ipsilateral channels, it is important to investigate which channels are driving this change. **Figure 4a** and **Figure 4b** show the change in log-transformed power in the alpha band over the baseline window of −250ms to −50ms before saccade onset for both the contralateral and ipsilateral channels. The relative change in alpha power shown in **Figure 3b** and **Figure 3c** is driven by an increase in ipsilateral power after saccade onset for both the Posner and neurofeedback tasks. Further there is a clear bilateral decrease in power that occurs just before the saccade onset in the Posner task, and at approximately 100ms after saccade onset in the neurofeedback task continuing past 400ms after saccade onset for both tasks. The topography of alpha activity from leftward minus rightward saccades indicates a posterior lateralisation after saccade onset **Figure 4c** and **Figure 4d**. This posterior lateralisation is similar for both the Posner and neurofeedback tasks but extends to the 200ms to 400ms period for the neurofeedback task only. In the current study, saccades were taken with visual stimuli present on the screen, and while participants were actively performing cognitive tasks. While this contrasts with the conditions reported in Liu et al., 2023, we report a replication indicating that these results are robust across different tasks and visual stimuli conditions.

**Figure 3:**
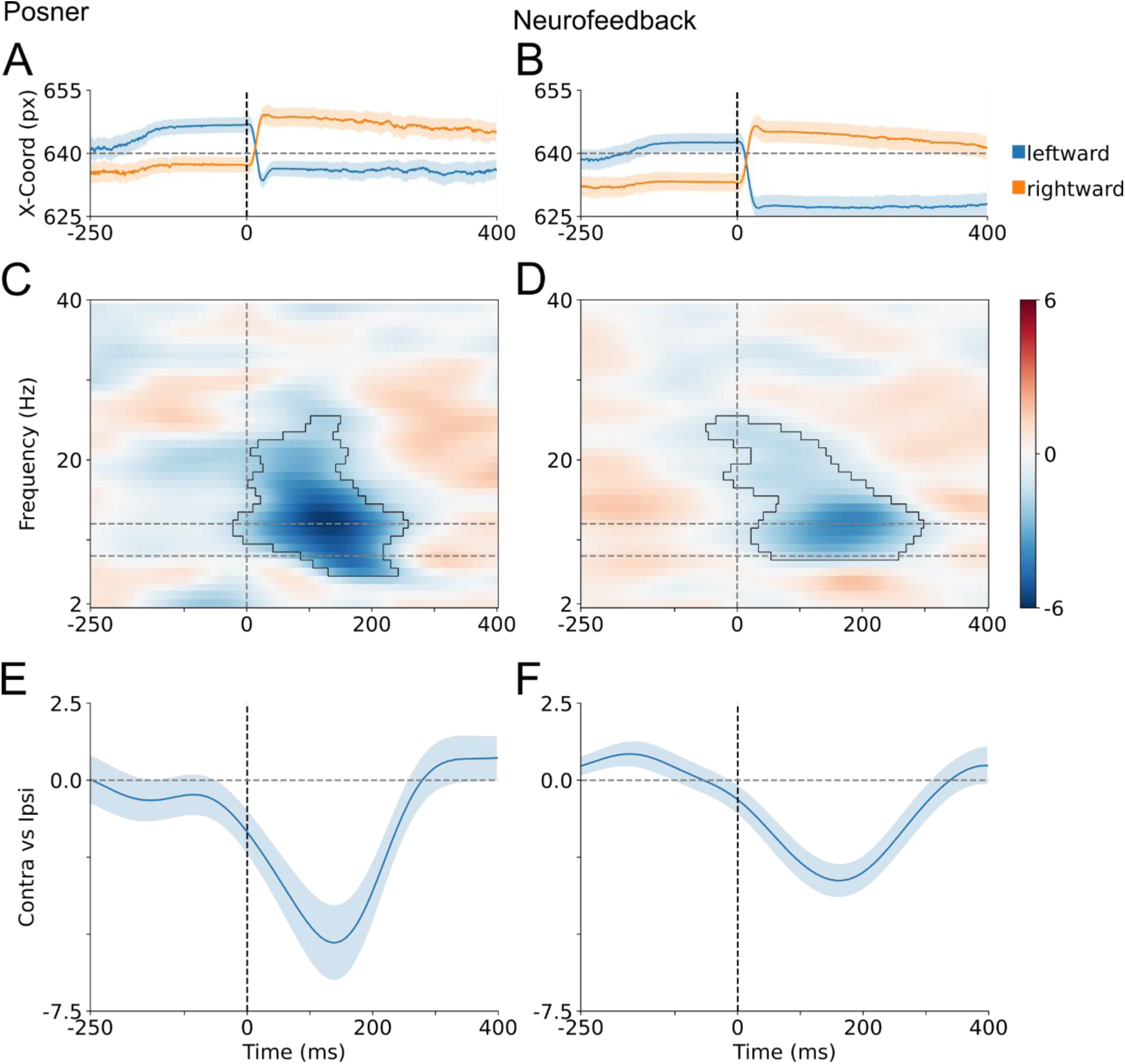
Saccade lateralisation results for both the Posner and neurofeedback tasks with saccade onset at 0ms. **A and B**) Saccade eye trace averaged over participants with centre of the screen located at 640 pixels represented by horizontal dashed line. The shaded area represents standard error of the mean. **C and D**) Time frequency plots showing difference between posterior contralateral and ipsilateral electrodes (PO7 and PO8) i.e. (contralateral – ipsilateral)/(contralateral + ipsilateral) relevant to saccade direction. Outlined areas show significant clusters and dashed horizontal lines represent alpha band (8-12Hz). **E and F**) Mean time course of alpha band for the contralateral and ipsilateral electrodes, relative to saccade direction. Shaded areas represent the standard error of the mean.

**Figure 4:**
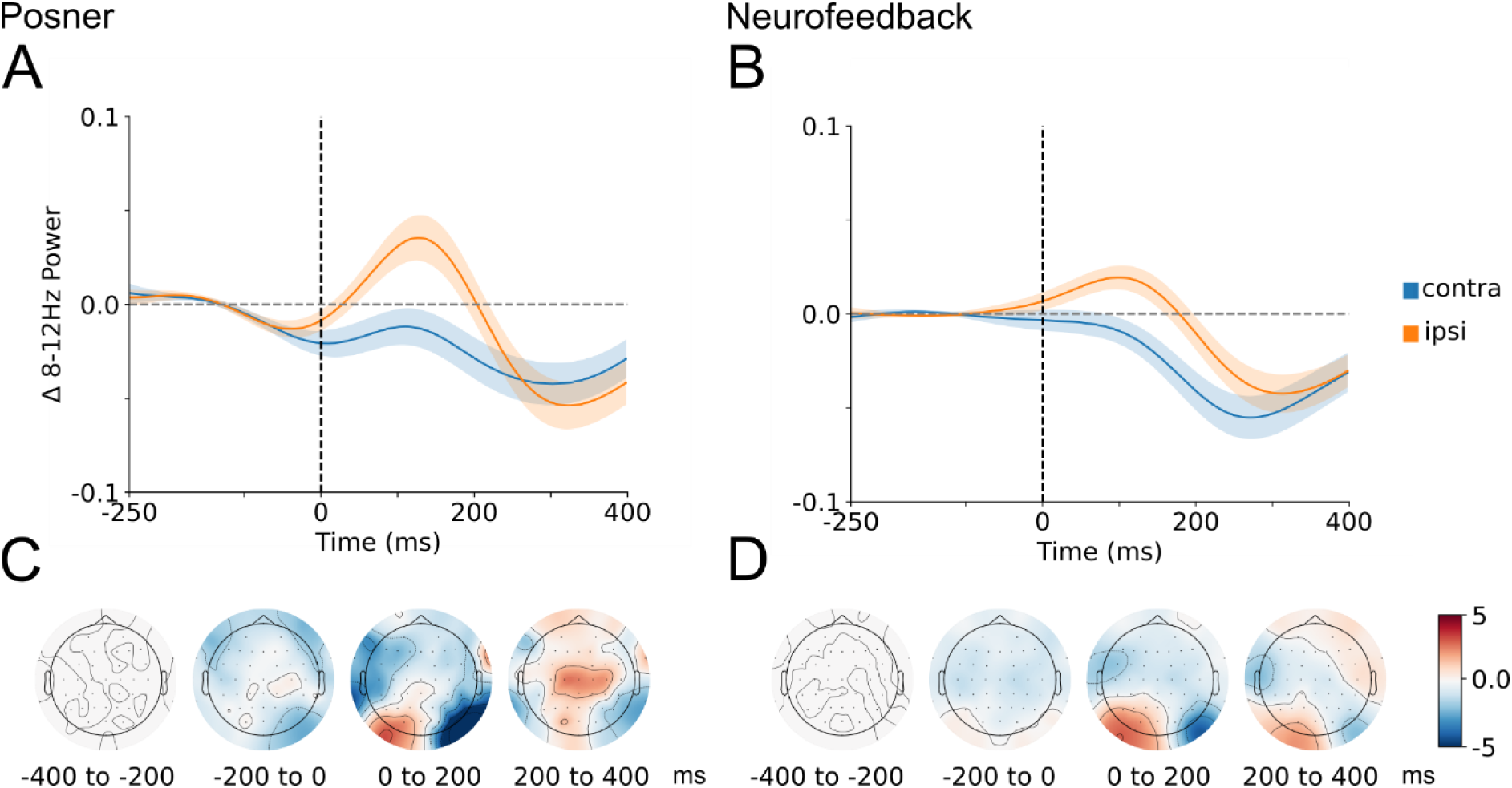
A and. **B**) Mean time course of alpha band power (8-12Hz) for the contralateral and ipsilateral electrodes showing change in log transformed power over baseline period of −250ms to −50ms before saccade onset. **C and D)** Time course of alpha band topography (left-right)/(left+right).

### 3.2 Effect of Visual Stimuli on EEG alpha lateralisation

A 2×2 repeated measures ANOVA comparing the spectral power between saccade groups (amplitude: small/large vs proximity: close/far) revealed no significant main effects and no interactions of saccade amplitude or saccade proximity for the Posner task. However, for the neurofeedback task, a significant main effect of saccade proximity was found with a significant cluster in the alpha band (cluster p-value = 0.04) (**Figure 5a**). No significant main effect of saccade amplitude and no interactions were observed for the neurofeedback task.

**Figure 5:**
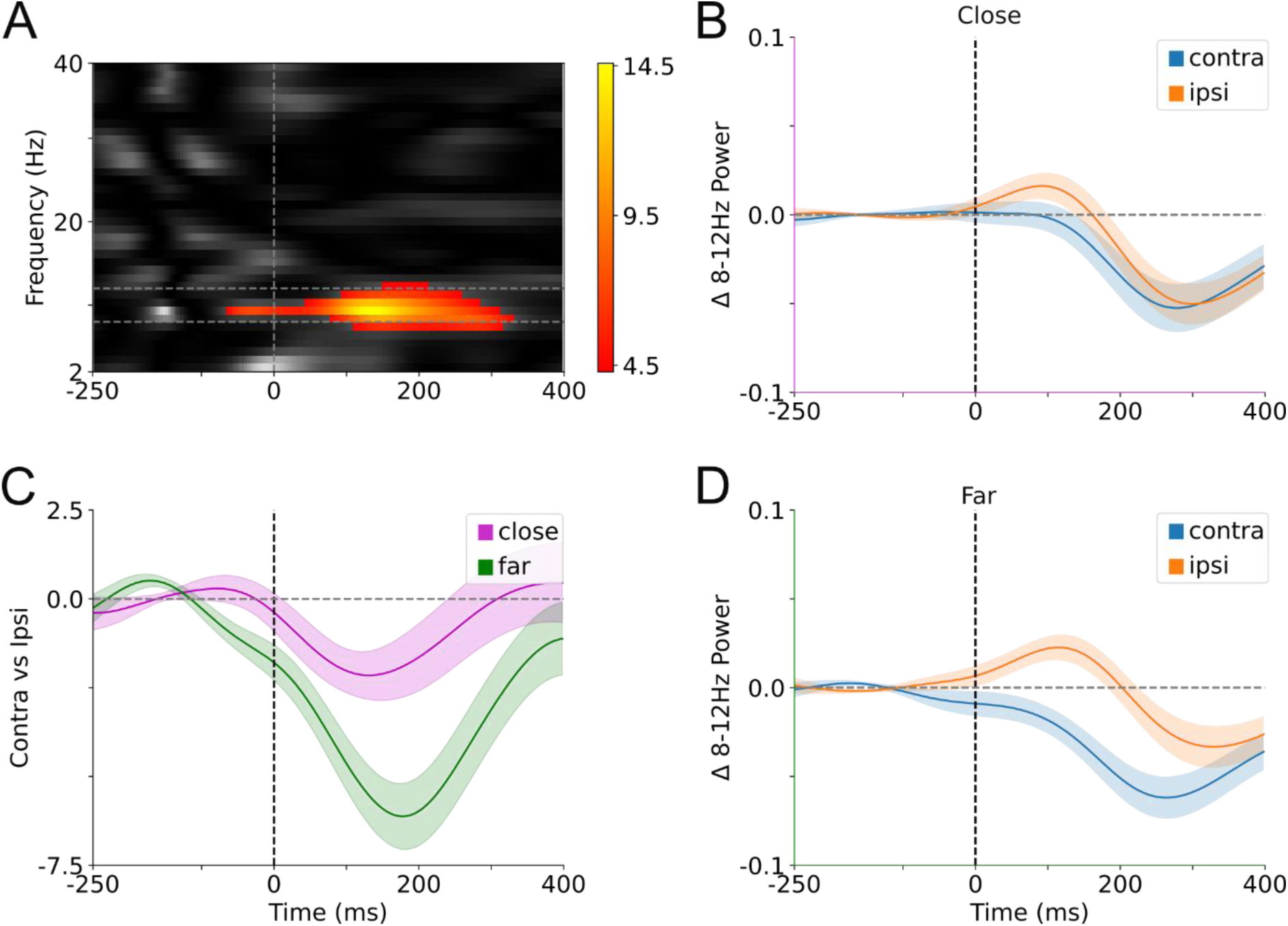
Main effect of saccade proximity to lateralised visual stimuli for the neurofeedback task. **A**) Time-frequency plot showing f-statistics with a significant cluster within the alpha band from approximately 50ms to 350ms after the saccade onset. **B**) Mean time course of contralateral and ipsilateral alpha power for saccades ending close to the visual stimuli. **C**) Mean time course of close and far alpha band lateralisation calculated over participants. Shaded area represents the standard error of the mean. Time courses are baselined by subtracting the mean of a baseline window from −0.25 to −0.5ms before saccade onset **D**) Mean time course of contralateral and ipsilateral alpha power for saccades ending far from the visual stimuli.

Investigating the main effect of saccade proximity in the neurofeedback task further, **Figure 5c** shows that saccades ending further from the lateralised stimuli elicited relatively larger lateralisation in the alpha band compared to those that ended closer to the lateralised stimuli. **Figure 5b** and **Figure 5d** show that this difference in lateralisation is due to both lower alpha power in the contralateral electrode and higher power in the ipsilateral electrode in the far saccade condition compared to the close saccade condition.

### 3.3 Saccade Distributions

Although it is not always the case, a saccade will often be larger to land close to the lateralised stimuli. Therefore, by design there was some overlap between the saccade groups. However, saccades do not always start precisely from the central fixation, thus the ‘close’ saccade group does not completely overlap with the ‘large’ group. **Table 2** shows the mean amplitudes of the saccades for the amplitude and proximity saccade groups in the Posner and neurofeedback tasks. The difference in mean amplitude for the close and far group is 2.85 times greater than the difference in mean amplitude for the large and small group for the Posner task and 3.43 times greater in the neurofeedback task.

**Table 2.** Saccade group mean amplitude.

|  | Posner | Neurofeedback |
| --- | --- | --- |
| Close | 0.86(0.41) | 1.00(0.51) |
| Far | 0.66(0.22) | 0.77(0.27) |
| Large | 1.01(0.47) | 1.28(0.55) |
| Small | 0.44(0.16) | 0.49(0.22) |

*Note: The mean(standard deviation) represents the grand average of saccade amplitudes for each saccade group. The units are degrees of visual angle.*

## 4 Discussion

In this study we observe a transient saccade-locked alpha lateralisation modulated by the proximity of saccade end-point location to on screen visual stimuli. Specifically, saccades which end close to on-screen visual stimuli are accompanied by weaker posterior alpha lateralisation (i.e. less increase in ipsilateral alpha power and less decrease in contralateral alpha power) compared to saccades that land further from visual stimuli during a covert spatial attention-based neurofeedback task. When considering all saccades together, we show that regardless of saccade size and endpoint location, posterior alpha power lateralisation post-saccade is present during active periods of two different tasks: a Posner reaction time task and a covert-spatial-attention based neurofeedback task. This lateralisation is driven by an increase in ipsilateral alpha power relative to contralateral alpha power and lasts for approximately 250-300ms post-saccade, replicating and extending previous research (Liu et al., 2023).

### 4.1 Saccade Induced Transient Alpha Lateralisation

When considering all saccades combined for each task, we report a transient, posterior lateralisation in the alpha band after saccade onset, replicating the results presented in Liu et al., 2023. We add to these results by analysing the EEG response to saccades obtained during two separate active cognitive tasks with visual stimuli present on the screen. The replication in the presence of the visual stimuli indicates that a large component of the reported posterior alpha is not affected by the specific task or visual context. This observation is in line with recent arguments proposing a tight coupling between lateralised posterior alpha power and the oculomotor system, suggesting this phenomenon is potentially the result of oculomotor planning and execution (Balestrieri et al., 2024; Kornrumpf et al., 2017; Liu et al., 2022; Popov et al., 2021).

Alternatively, saccade-locked alpha lateralisation could be visual in nature. Visual input is partially suppressed during the peri-saccadic interval. In vitro studies have shown that suppression starts as early as the retina (Idrees et al., 2020). Thus, when suppression abruptly stops at the end of a saccade, a new flow of visual input can impinge on the retina. This occurs irrespectively as to whether the pre-saccadic and the post-saccadic inputs are the same and indeed happens also when eye movements are performed over a uniform grey screen. In this case the crucial comparison is between the suppressed visual input *during* a saccade, and the un-suppressed *post-saccadic* visual input. In this alternative account, the main driver of saccade-locked alpha lateralisation would be visual in nature, and one would expect saccade-locked alpha lateralisation to manifest itself regardless of task and visual context. It is crucial to note that in this case a lateralised signal would emerge from the different visual input between contralateral and ipsilateral visual fields, for example due to the uneven visual stimulation caused by the proximity of screen borders or other visual stimuli.

### 4.2 Saccade Lateralisation and Proximity

The main effect of proximity to visual stimuli in the lateralisation of alpha power lends support for a role of visual afference in modulating posterior alpha lateralisation after a saccade. In addition to the change in retinal input following saccadic suppression, it has previously been demonstrated that alpha desynchronisation is spatially tuned to visual stimuli and may predict alpha lateralisation responses during attention cueing experiments (Yuasa et al., 2023). Therefore, if alpha lateralisation can depend on the location of visual stimuli on the fovea, a centrally presented stimulus would result in a more balanced distribution of posterior alpha power across hemispheres, i.e., weaker lateralisation. In our case, saccades landing closer to visual stimuli would place those stimuli more central in the visual field than those landing further away. This would explain the weaker lateralisation that we see with the ‘close’ saccades compared to the ‘far’. Liu et al., 2023 highlight that the alpha response to new visual stimuli can be slow, peaking around 500ms (e.g. Boettcher et al., 2021) compared to the 150-200ms that we report. However, there are also studies that report stimulus-evoked lateralised alpha in a shorter time window, from as low as 100ms after cue onset (Arana et al., 2022; Balestrieri et al., 2024). Therefore, the difference in alpha lateralisation due to shifting visual stimuli on the fovea cannot be ruled out.

Another, although not mutually-exclusive, explanation for the difference in alpha lateralisation due to saccade proximity to visual stimuli is due to inhibition of irrelevant visual inputs (Foxe & Snyder, 2011; Jensen, 2023; Jensen & Mazaheri, 2010). It is argued that alpha oscillations play a role in gating visual stimuli by inhibition, i.e. increased alpha may correspond to a suppression of the contralateral visual field containing non-relevant visual information. It is possible that the increase in ipsilateral power we observe for saccades that land further from visual stimuli is the result of higher suppression of the ‘undesired’ target. As there are always two lateralised stimuli present on the screen, when a saccade lands close to one it is inherently further away from the other. Additionally, a saccade landing far from the stimulus it was directed to will result in more of the opposite stimulus remaining in the visual field. For example, a leftward directed saccade landing somewhere in the middle of the screen (far from the left stimuli) will result in more of the right (undesired) stimulus present in the visual field leading to more inhibition of the right stimulus.

### 4.3 Saccade Amplitude Does Not Modulate Alpha Lateralisation

Our results do not show a main effect of saccade amplitude in either the Posner or Neurofeedback tasks. There is some overlap of saccades between the proximity and amplitude groups as it requires a larger saccade to shift from central fixation to the lateralised stimuli resulting in more ‘close’ saccades in the ‘large’ saccade group. However, for the neurofeedback task, the difference in the mean amplitude between the ‘small’ and ‘large’ group is 3.4 times larger than the difference in mean amplitude between the ‘close’ and ‘far’ groups. Therefore, if the effect reported was driven by saccade amplitude, we would expect to see a larger difference in lateralisation in the amplitude group. Instead, we only find a main effect of proximity suggesting that the amplitude of saccades does not impact the subsequent EEG lateralisation. This is in line with previous findings suggesting that the amplitude of saccades result in similar evoked responses (Dimigen et al., 2009), downplaying a potential contribution of saccade-locked corollary discharge on alpha lateralisation.

### 4.4 Only Neurofeedback Task Shows Modulation of alpha Lateralisation

In the current study we only find a main effect of saccade proximity to visual stimuli in the neurofeedback task. This could firstly be explained by an issue of statistical power as there are over three times as many saccades used in the analysis for the neurofeedback task compared to the Posner task. This is because the saccades in the neurofeedback task were taken from a two-minute window, whereas the saccades in the Posner task were taken from a period of less than two seconds. Furthermore, the longer neurofeedback tasks means that saccades are taken from a much more stable period compared to the Posner task, potentially removing noise in the EEG recordings.

Another explanation for the lack of effect in the Posner task is the difference in visual stimulation and cognitive tasks. The neurofeedback task is characterised by a dynamically changing stimulus on the left and a static (black) stimulus on the right. In contrast, the Posner task has static blue stimuli on both sides. Even though the Posner and neurofeeback tasks are spatially laid out in an almost identical way, the neurofeedback task forces participants to adjust covert attention to the left stimulus while the flashing stimulus simultaneously elicits exogenous attention. In contrast, the Posner task, implicitly directs endogenous attention, which can be left, right, and neutral. As fixational eye movements are heavily linked to spatial attention (Casteau & Smith, 2020; Engbert & Kliegl, 2003; Hafed et al., 2015; Hafed & Clark, 2002), it is possible that the differences in the task could affect the planning, execution, and suppression of saccades despite similar saccade dynamics between them. In turn, a potential oculomotor account of alpha lateralisation (Popov et al., 2021) would predict a difference between the two tasks.

Further, Covert spatial attention is strongly linked to lateralised alpha power (Kornrumpf et al., 2017; Peylo et al., 2021; Rihs et al., 2007; Sauseng et al., 2005; Thut et al., 2006). Thus, the underlying covert spatial attention task, particularly for neurofeedback, could potentially be a driver for the lateralised alpha that we see post-saccade. However, we only find an effect specific to the saccade proximity group and not the saccade amplitude group. If our results were driven by covert spatial attention, we would expect comparable lateralisation in the saccade groups, particularly as there is some overlap between the proximity and amplitude groups.

### 4.5 Limitations

One limitation of this study is that the neurofeedback task required participants to perform leftward covert spatial attention constantly during a trial while, at the same time, the left lateralised visual stimulus was dynamically changing colour. Therefore, the reported effects are in the presence of sustained lateralisation due to both events. We control for this by splitting saccades based on their amplitude in addition to proximity to visual stimuli. As we only observe an effect due to proximity, we conclude that this is above any underlying, systematic effects such as continuous lateralised attention. Nonetheless, as all saccades are taken from active attention tasks, we cannot rule out a component of lateralised attention, potentially confounding the results of both saccade groups. Further investigations would be required with a control saccade group taken from a similar time duration as the task trials but from a rest period (undertaking no specific task) with no visual stimuli present on the screen to eliminate this possibility.

## 5 Conclusion

We observe reliable post-saccadic lateralisation of posterior alpha power across different tasks. This lateralisation can be modulated by the landing proximity of the saccades to on-screen visual stimuli but not by saccade amplitude, indicating a central role of either retinal displacement of the visual stimulus while the eyes are moving, or the influence of immediate post-saccadic visual afference. Our results highlight both the importance of recording eye movements during EEG experiments investigating alpha lateralisation to control for oculomotor confounds, as well as the structure and properties of the visual stimuli present during the tasks.

## 6 Data and Code Availability

Raw data in BIDS format in addition to scripts for analysis and visualisation are available on the Open Science Framework (OSF) (Turner et al., 2022). Note, the contained MATLAB script *groupSubjects_bids.m* for identifying saccades depends on code from the Niehorster et al., 2015 adaption of the Nyström & Holmqvist, 2010 algorithm. This code can be found here: https://github.com/dcnieho/NystromHolmqvist2010/tree/master and has also been uploaded alongside our scripts on OSF.

## 7 Author Contributions

C.T.: Conceptualisation, Methodology, Software, Formal analysis, Investigation, Data Curation, Writing - Original Draft, Writing - Review & Editing, Visualisation, Project administration, Funding acquisition

A.V. Supervision, Conceptualisation, Writing - Review & Editing, Funding acquisition

G. L. Supervision, Conceptualisation, Resources, Writing - Review & Editing, Funding acquisition

A.F. Supervision, Conceptualisation, Software, Resources, Writing - Review & Editing, Funding acquisition

## 8 Funding

C.T. was supported by an EPSRC studentship (EP/T517896/1)

A.V. EPSRC studentship (EP/T517896/1)

G.L. was supported by the Wellcome Trust [209209/Z/17/Z].

A.F. was supported by a grant from the Biotechnology and Biology Research Council (BBSRC, grant number: BB/S006605/1) and the Bial Foundation (Bial Foundation Grants Programme; Grant id: A-29315, number: 203/2020, grant edition: G-15516).

